# *Cis*-regulatory elements orchestrate phase-specific effector gene expression in *Ustilago maydis*

**DOI:** 10.64898/2026.03.26.714514

**Authors:** Georgios Saridis, Janina Werner, Katharina Stein, Luyao Huang, Ute Meyer, Jan Muelhoefer, Namarta C. Singh, Gunther Doehlemann

**Affiliations:** Institute for Plant Sciences, University of Cologne, Cologne, Germany and Cluster of Excellence on Plant Sciences (CEPLAS), University of Cologne, Cologne, Germany

**Author notes:** Equal contribution (Georgios Saridis, Janina Werner).

## Abstract

Plant pathogenic fungi secrete small proteins, termed effectors, to reprogram host metabolism and suppress immune responses during infection. Although transcriptional waves of effector expression have been described in several pathosystems, the *cis*-regulatory elements encoding infection-stage specificity remain largely unknown. Here, we investigate the temporal regulation of effector genes in the biotrophic smut fungus *Ustilago maydis*, a model organism for fungal plant pathogenesis. By integrating transcriptome reanalysis with comparative promoter motif enrichment across biotrophic fungi, we identify distinct promoter motifs associated with defined infection phases. In *U. maydis*, three candidate *cis*-regulatory elements correlate with early, proliferative, and late infection stages, respectively. Positional enrichment relative to transcription start sites supports their regulatory relevance. Functional promoter mutagenesis demonstrates that the early-phase motif GTGGG significantly contributes to effector gene expression *in planta* and is sufficient to drive stage-restricted gene expression in synthetic minimal promoters.

Collectively, our findings demonstrate that temporal deployment of the effector repertoire is at least partially encoded at the promoter level. The identified *cis*-regulatory elements provide a framework for dissecting transcriptional control during biotrophic infection and offer tools for infection-stage-specific gene expression in synthetic biology applications.

## Introduction

To effectively fulfill their infection cycle, plant pathogenic fungi secrete small proteins, called effectors. Effectors have diverse roles at the apoplastic space between the interacting parties, or when inserted in the host cells, mainly manipulating the host metabolism and/or suppressing host defenses (Thomma et al., 2011). Considering that these proteins are valuable for host colonization, they are frequently recognized and targeted by the host immune system, ushering a process of selection pressure best described as the arms race (Ingle et al., 2006). This competitive evolutionary mechanism renders effectors as vital, fast-evolving proteins with a fundamental potential as study subjects for the better understanding of pathogen-host interactions, as well as the development of pathogen-negating agricultural approaches (Fouché et al., 2018). Rapid effector diversification raises the question whether regulatory logic may be more conserved than effector sequence itself.

Smut fungi are biotrophic plant pathogens of the Ustilaginales order, which infect a wide range of plants, from monocots to dicots, including economically important cereal crops, such as barley, wheat and maize. The corn smut *Ustilago maydis* is an established model system for biotrophic fungal genetics, being ranked amongst the top 10 of the most scientifically significant fungal pathogens (Dean et al., 2012; Matei & Doehlemann, 2016). It has two forms, namely the saprophytic haploid yeast-like form, which does not cause any disease and can be grown in artificial media, and the biotrophic dikaryotic filamentous form, which is generated by the fusion of two compatible haploid cells (Kämper et al., 2006; Kruzel & Hull, 2010). The infection of *U. maydis* cycle comprises (i) penetration, (ii) *in planta* proliferation, and (iii) tumor formation and sporulation, the latter being the typical smut phenotype (Banuett & Herskowitz, 1996; Brefort et al., 2009). It is therefore highly likely that *U. maydis* infection of maize relies on the deployment of phase-specific effectors.

Many *U. maydis* effectors have been studied and characterized for their function and mode of action. Additionally, a number of transcriptional regulators have been identified and characterized, separating the activation of its effectome repertoire into two waves (Djamei et al., 2023). The early effectome activation is massively linked to *U. maydis* cell fusion and filamentation, which leads to the activation of the *b* mating-type locus genes and the production of the heterodimeric bE/bW homeodomain transcription factor (Gillissen et al., 1992; Kämper et al., 1995). This master regulator of the early pathogenic development activates in turn a group of effector genes as well as the zinc finger transcription factor Rbf1 and its target transcription factors Hdp2 (homeobox), Biz1 (C_2_H_2_ zinc finger), and Mzr1 (C_2_H_2_ zinc finger) (Flor-Parra et al., 2006; Heimel et al., 2010; Zheng et al., 2008). Uncharacterized transcription factors correlated with early effectome expression include proteins containing bHLH, TEA/ATTS, HMG, CCHC, and winged-helix domains (Lanver et al., 2018). Upon *in planta* proliferation and hyphal development, Zfp1, a Zn_2_Cys_6_ transcription factor, has been connected with the regulation of 112 effectors and 50 transcription factors, including Ros1 (WOPR), which successively activates the late effectome while simultaneously attenuating the wave of early effectors during sporulation and tumor formation (Cheung et al., 2021; Tollot et al., 2016). Additionally, the previously characterized transcription factors Nlt1 (APSES), Fox1 (forkhead) as well as novel proteins including ones containing Zn_2_Cys_6_, bHLH, C_2_H_2_, and winged-helix domains, have been associated with effector gene expression at the late stages of infection (Lanver et al., 2017, 2018; Zahiri et al., 2010). Despite detailed knowledge of the transcription factor hierarchy controlling pathogenic development, the *cis*-regulatory elements that encode infection-stage-specific effector expression remain unidentified. Here, we systematically identify promoter motifs associated with infection-phase-specific effector expression in *U. maydis* using comparative genomics, transcriptome reanalysis, and functional promoter validation. By integrating evolutionary and stage-specific analyses, we ask whether temporal effector deployment is encoded at the promoter level.

## Results

### Comparative identification of effector-associated promoter motifs in biotrophic fungi

To identify putative *cis*-regulatory elements enriched in the promoters of fungal effector genes, the genomes of different smuts, rusts and powdery-mildews were analyzed (**Figure 1A**). Protein fasta files from nine smut fungi (*Ustilago maydis, U. bromivora, U. tritici, U. nuda, U. loliicola, U. hordei, Sporisorium reilianum, S. scitamineum, Melanopsychium pennsylvanicum*), five rust fungi (*Puccinia sorghi, P. striiformis f. sp. tritici, P. triticina, P. graminis, Melampsora larici-populina*) as well as five powdery-mildew fungi (*Blumeria hordei, B. graminis f. sp. tritici, Erysiphe necator, E. neolycopersici, Golovinomyces cichoracearum*) were subjected to identify orthologous genes among the tested fungal species. Based on the orthologous analysis the species tree was constructed using *Saccharomyces cerevisiae* as a general outgroup, as well as *Testicularia cyperi* as outgroup for the smut clade (**Figure 1B**). Simultaneously, for each species, those files were used as input to predict their total effectome. The Phobius tool was initially used to predict transmembrane domains in protein sequences, followed by signaling peptide via SignalP. Proteins with transmembrane domains and without a signaling peptide were filtered out, while the remaining sequences were labelled as secretome and were thereafter used for an analysis with EffectorP to predict the total effectome, leading to effectome sizes spanning from 157 in *U. loliicola* to 1266 in *P. graminis* (**Table S1**). The 500 bp promoter sequences from all predicted effector genes for each species were used to identify enriched motifs using the tool XSTREME from the MEME Suite. The predicted motifs which passed the thresholds (*P*-value < 0.05 and enrichment sites > 10%) were compared with each other using STAMP. The smut *M. pennsylvanicum* and the powdery-mildews *E. necator, G. cichoracearum* and *E. neolycopersici* showed no significant motif enrichment. Notably, one motif with consensus sequence GTGGG was specifically enriched in the effectome promoters of all tested smuts but not in rusts and powdery-mildews (**Figure S1**). Two rust-specific enrichments were identified with consensus sequences KCGA and TGCK, while further consensus motifs derived from either smuts and powdery-mildews, such as ATGAA, powdery-mildews and rusts, such as CGTG, or smuts and rusts, including CWGTR and CGGCS (**Figure S1**).

**Figure 1.**
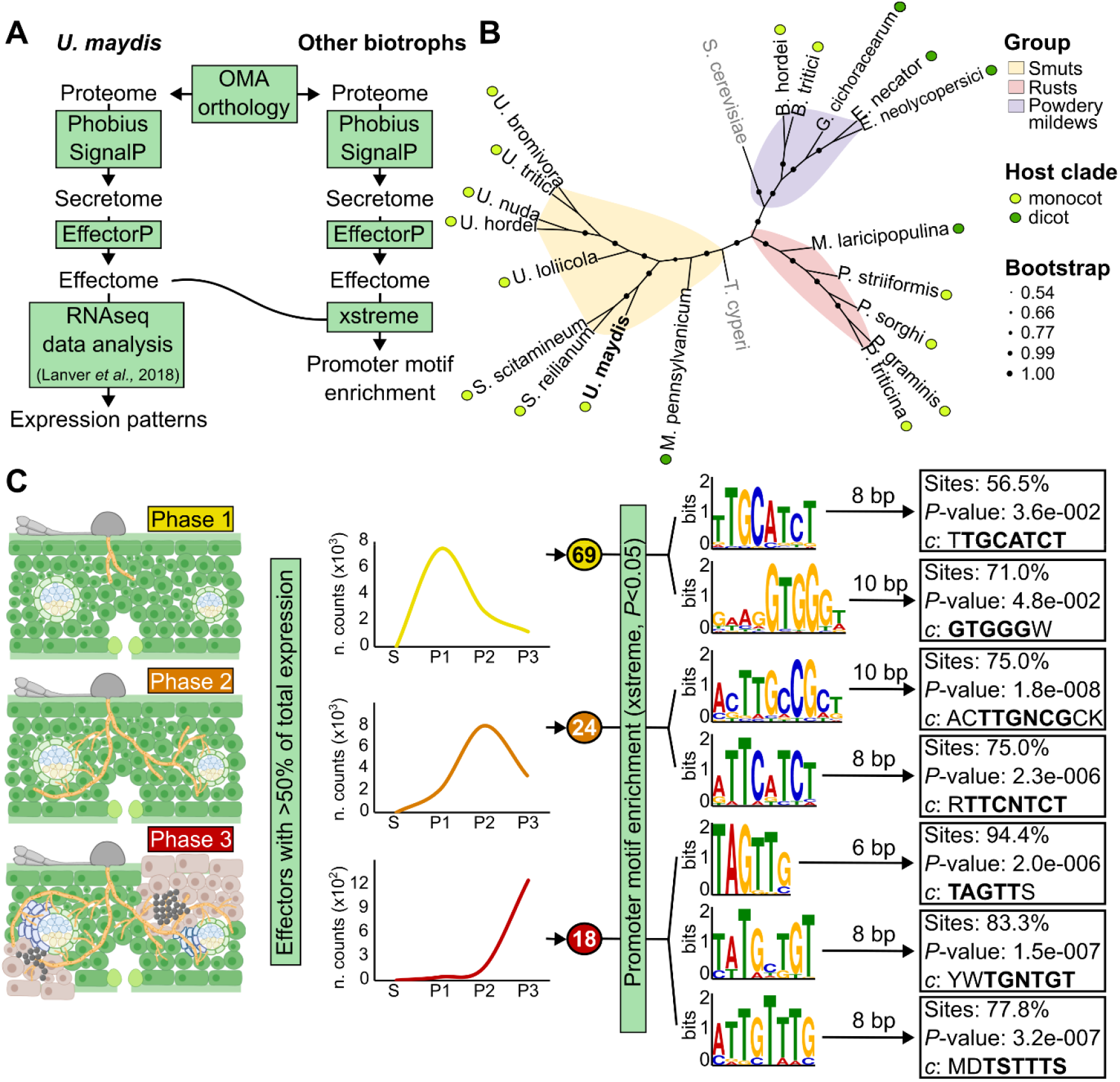
Identification of putative phase specific motifs in *Ustilago maydis* effectome promoters. **(A)** Workflow of the bioinformatic analysis for the prediction of putative effectomes in the tested species used for the promoter enrichment analysis. **(B)** Phylogenetic tree illustrating the evolution of the species used in this study. **(C)** Selection of phase specific effector genes in *U. maydis* and promoter motif enrichment. Motifs with > 55% enriched sites as well as significant of enrichment *p*-value<0.05 are shown here and were used for the follow-up analysis. Bold letters in the motif consensus (labelled as *c*) indicate a putative core sequence. n. counts: normalized counts in counts per million.

### Infection-phase-specific clustering of *U. maydis* effectors and motif enrichment identifies candidate *cis*-regulatory elements

To dissect phase-specific effector regulation in *U. maydis*, we reanalyzed RNA-seq data generated by Lanver et al. (2018) and grouped infection into three biologically defined stages: (i) Phase 1 (0.5–2 dpi; penetration and early colonization), (ii) Phase 2 (4-6 dpi; hyphal proliferation), and (iii) Phase 3 (8–12 dpi; tumor formation and sporulation) (**Figure 1C, Table S2**). Effector genes were assigned to a phase if more than 50% of their cumulative expression occurred within that interval. This classification yielded 69 Phase 1-, 24 Phase 2-, and 18 Phase 3-enriched effectors **(Figure 1C, Table S3**), consistent with temporally structured transcriptional waves during infection (**Figure 1C, Table S3**).

Promoter regions (500 bp upstream) of phase-enriched effectors were subjected to motif enrichment analysis using XSTREME. Seven significantly enriched motifs (P < 0.05; enrichment sites >55%) were identified across the three infection phases. Two motifs were associated with Phase 1 (TGCATCT [P1-1] and GTGGG [P1-2]), two with Phase 2 (TTGNCG [P2-1] and TTCNTCT [P2-2]), and three with Phase 3 (TAGTT [P3-1], TGNTGT [P3-2], and TSTTTS [P3-3]) (**Figure 1C**). Among these, GTGGG (Phase 1), TTGNCG (Phase 2), and TSTTTS (Phase 3) displayed the strongest phase-restricted enrichment patterns and were prioritized for further analysis.

### Intra- and inter-clade specificity of the obtained motifs

To assess motif conservation and specificity, we calculated enrichment of each motif in effector versus non-effector promoter sets across all tested species. For this, the enrichment of every motif in each species effectome versus non-effectome (background) promoters was calculated. For the motif TGCATCT [P1-1], 6 out the 9 tested smuts showed enrichments in the effectome, with *U. maydis* demonstrating the highest enrichment, while for 3 of those, namely *S. scitamineum, S. reilianum* and *M. pennsylvanicum*, the enrichment was towards the background (**Figure 2A**). No significant inter-clade differences were observed. For the other motif of Phase 1, GTGGG [P1-2], an inter-clade difference was present with all smut motifs clustering closely and demonstrating a strong preference of this motif towards smut effectome promoters (**Figure 2A**). Motif TTGNCG [P1-2] revealed an overall negative enrichment (towards the background) for all fungal clades. However, individual species (*U. maydis, U. bromivora, S. scitamineum, P. sorghi, P. graminis, B. hordei, Golovinomyces cichoracearum* showed effectome enrichment of the P2-1 motif. Motif TTCNTCT [P2-2] had a powdery-mildew-specific enrichment, with all its tested species demonstrating effectome enrichment, while from smuts, only 4 out of 9 showed a similar pattern **(Figure 2A**). Amongst the smuts, *U. maydis, U. loliicola, S. reilianum* and *M. pennsylvanicum, U. maydis* had again the strongest enrichment. Motifs TAGTT [P3-1] and TGNTGT [P3-2] demonstrated very similar patterns with a preference of those motifs towards smut effectomes, with only exceptions *U. hordei* for motif P3-1 and *U. tritici* for motif P3-2 (**Figure 2A**). In both cases, the highest enrichment was observed for *U. maydis*. Finally, motif TSTTTS [P3-3] showed an overall enrichment towards the background for smuts and rusts and a slight enrichment towards the effectome of powdery-mildews (**Figure 2A**). Nevertheless, *U. maydis, S. reilianum* and *U. tritici* were the only smuts with an effectome enrichment.

**Figure 2.**
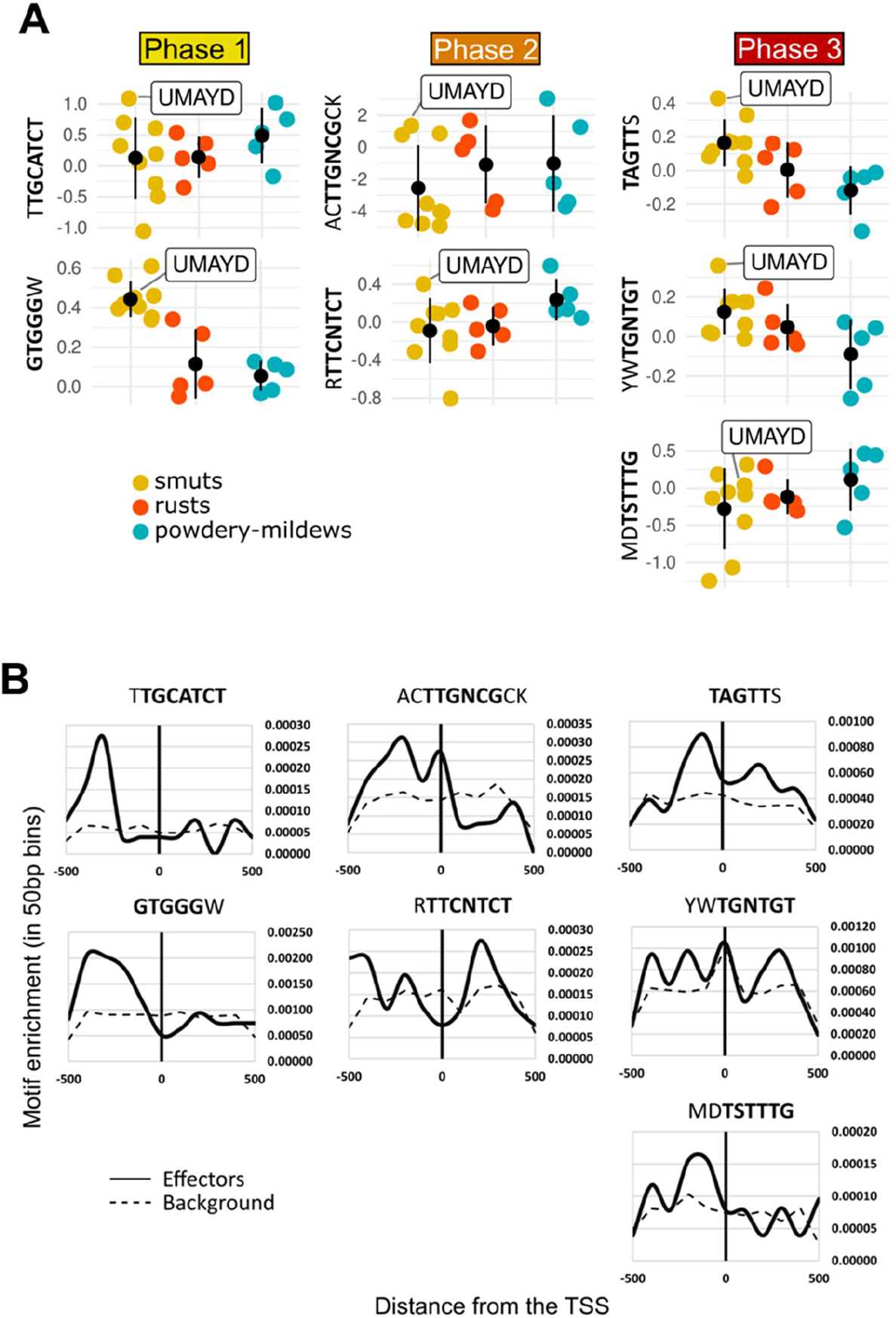
Specificity of motif enrichment within and among fungal clades and motif presence around the TSS in *U. maydis*. **(A)** Colored dots represent the enrichment of each motif in the effectome of a species versus the non-effectome. Black dots with error bars represent the average as well as standard deviation, of the motif enrichment within each fungal clade. Bold letters of the motifs indicate the putative core sequence. **(B)** Motif occurrence was calculated in all effector and non-effector (background) gene promoters in bins of 50 bp, 1 kb around the TSS. The average enrichment for each bin was then calculated and the enrichment curve was plotted. Bold letters of the motifs indicate the putative core sequence. Solid line: effectome, dashed line: non-effectome.

### Positional enrichment supports regulatory relevance

Next, the presence of each motif in a 1kb region, spanning around the transcription start site (TSS) for both the effectome and non-effectome background, was calculated, and their frequencies were plotted (**Figure 2B**). Several motifs showed non-random positional enrichment upstream of the TSS (Figure 3). GTGGG [P1-2] displayed a broad enrichment between 150–450 bp upstream of the TSS, consistent with promoter-proximal regulatory regions. TTGNCG [P2-1] showed enrichment across the 0–450 bp upstream region, whereas TSTTTS [P3-3] exhibited discrete upstream peaks. Some motifs also displayed downstream enrichment. These patterns may reflect alternative transcription start sites, regulatory elements within 5′ untranslated regions, or limitations of current gene annotations. Overall, the positional bias of multiple motifs supports their potential regulatory function. Additionally, we tested whether known transcription factor binding sites could potentially correlate with our motifs. For this, we subjected the consensus core sequences of each motif to an analysis with TomTom from the MEME Suite, using the JASPAR Fungal database. The P1-1 motif could be similar to an APSES-type binding site (*P*-value = 0.0011), while the P1-2 motif could be targeted by C_2_H_2_ zinc finger transcription factors (*P*-value = 0.0003). Additionally, the P2-1 motif might be resembling another C_2_H_2_ binding site (*P*-value = 0.0019), while the P2-2 motif is likely a binding site for a heat shock factor (*P*-value = 0.0023) (**Table S4**). Finally, the P3-1 motif is predicted as binding site for a homeodomain (*P*-value = 0.0154), the P3-2 motif for a bZIP (*P*-value = 0.0291) and the P3-3 motif for a forkhead/winged helix (*P*-value = 0.0082) transcription factor (**Table S4**).

**Figure 3.**
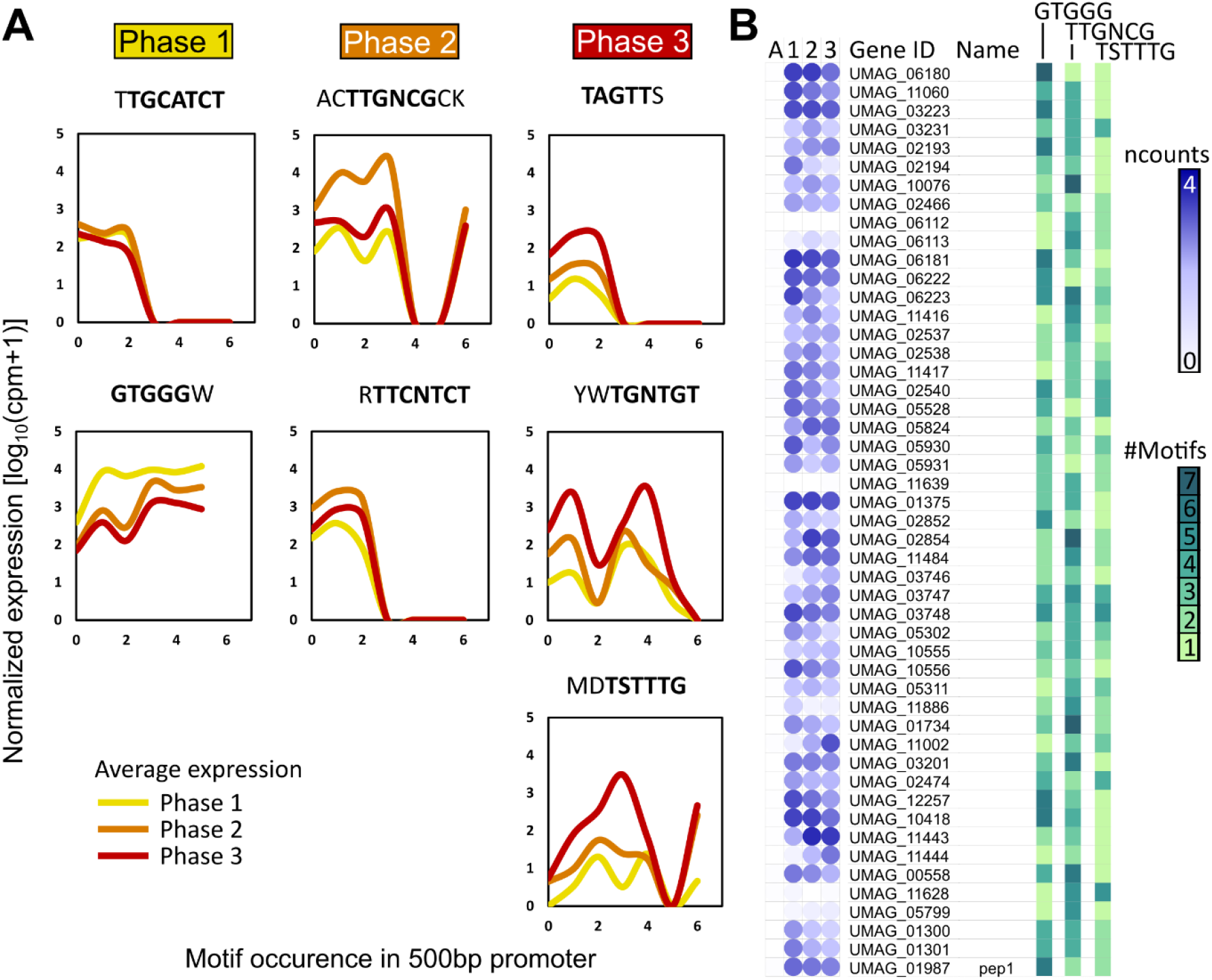
Estimation of the phase-specific activity of the motifs. **(A)** Correlation between the number of motifs in the promoters and the gene expression in each phase. The average expression value for each phase was calculated for all the genes with the same amount of each motif in their 500 bp promoter. Bold letters of the motifs indicate the putative core sequence. **(B)** Selected genes which contain all three of the candidate phase-specific motifs. Heatmap (circles) depicts normalized counts (ncounts, in log_10_(cpm+1)). The numbers of each motif are shown in the right panel.

### Motif occurrence correlates with phase-specific gene expression

To investigate the phase-specificity of the identified motifs, we performed a correlative analysis of the effectome gene expression and the motif occurrence in their promoters. We used the maximum expression potential of each effector gene and correlated that to the number of copies present in their promoters. GTGGG [P1-2] showed a positive association between motif presence and Phase 1 expression levels (**Figure 3A**). Promoters containing one or two GTGGG copies exhibited elevated early-stage expression compared to promoters lacking the motif (**Figure S2**). Therefore, we wanted to further dissect the possible influence of each motif to the phase-specific expression. To this, *U. maydis* effectors were split in groups based on the amount of each motif present in their promoters, ranging between 0-6 occurrences, and then the average expression of each group in each infection phase was calculated and plotted. Motif P1-1 (TGCATCT) showed highest expression with up to two copies but no phase specificity (**Figure 3A**). P2-1 (TTGNCG) enabled expression with 0–3 or 6 copies, with a strong Phase 2 preference at 1–3 copies. P2-2 (TTCNTCT) resembled P1-1, allowing expression only up to two copies without phase specificity. P3-1 (TAGTT) showed a similar pattern, with slight Phase 3 preference. P3-2 (TGNTGT) exhibited higher expression at 1 or 3–4 copies, with weak Phase 3 specificity at 1 or 4 copies. Finally, the motif TSTTTS [P3-3] had the clearest specificity towards expression at Phase 3, especially when there were 3-4 copies of this motif present in the promoters (**Figure 3A**).

The best performing motifs, namely GTGGG [P1-2], TTGNCG [P2-1] and TSTTTS), were used to identify the most fitting effector candidate for their follow-up experimental characterization. 49 effectors containing all three motifs in their 1 kb promoters were selected and their expression was determined using the re-analyzed RNAseq dataset from Lanver et al. (2018) (**Figure 3B**).

### Functional validation identifies GTGGG as an early infection regulatory module

To experimentally test whether enriched motifs directly influence effector gene expression, we performed promoter mutagenesis using the *pep1* promoter driving GFP expression at the *ip* locus. In our experimental setup, we incorporated mutations of the GTGGG [P1-2] motif, in positions -56, -160, -240, -300, -394, -584, 2 copies of the TTGNCG motif [P2-1] in positions -671 and -839, and 2 copies of the TSTTTS motif [P3-3] found in positions -720 and -885. To ensure the success of the proof-of-concept experiments, all mutations were based on the least probable nucleotide at the respective position, according to the confusion matrix of the identified motif (**Table S5**). We have selected the following time points: for Phase 1, 3 dpi; for Phase 2, 6 dpi; for Phase 3, 10 dpi.

Three fungal mutants were generated: One where the P1-2 motif was mutated, one where the P2-1 and P3-3 motifs were mutated and one where P1-2, P2-1 and P3-3 were mutated (mp_ALL_). The resulting *U. maydis* strains were inoculated into maize seedlings and compared with the wild-type p^*pep1*^-GFP strain where no motifs were mutated. Quantification of *gfp* levels at 3-, 6-, and 10-days post inoculation (dpi) revealed a significant reduction in *gfp* expression upon mutation of the Phase 1 motif or all motifs combined, whereas mutation of the phase 2 and 3 motif had no detectable effect (**Figure 4A,B**). These results demonstrate that the P1-2 motif contributes significantly to *pep1* promoter activity during early infection.

**Figure 4.**
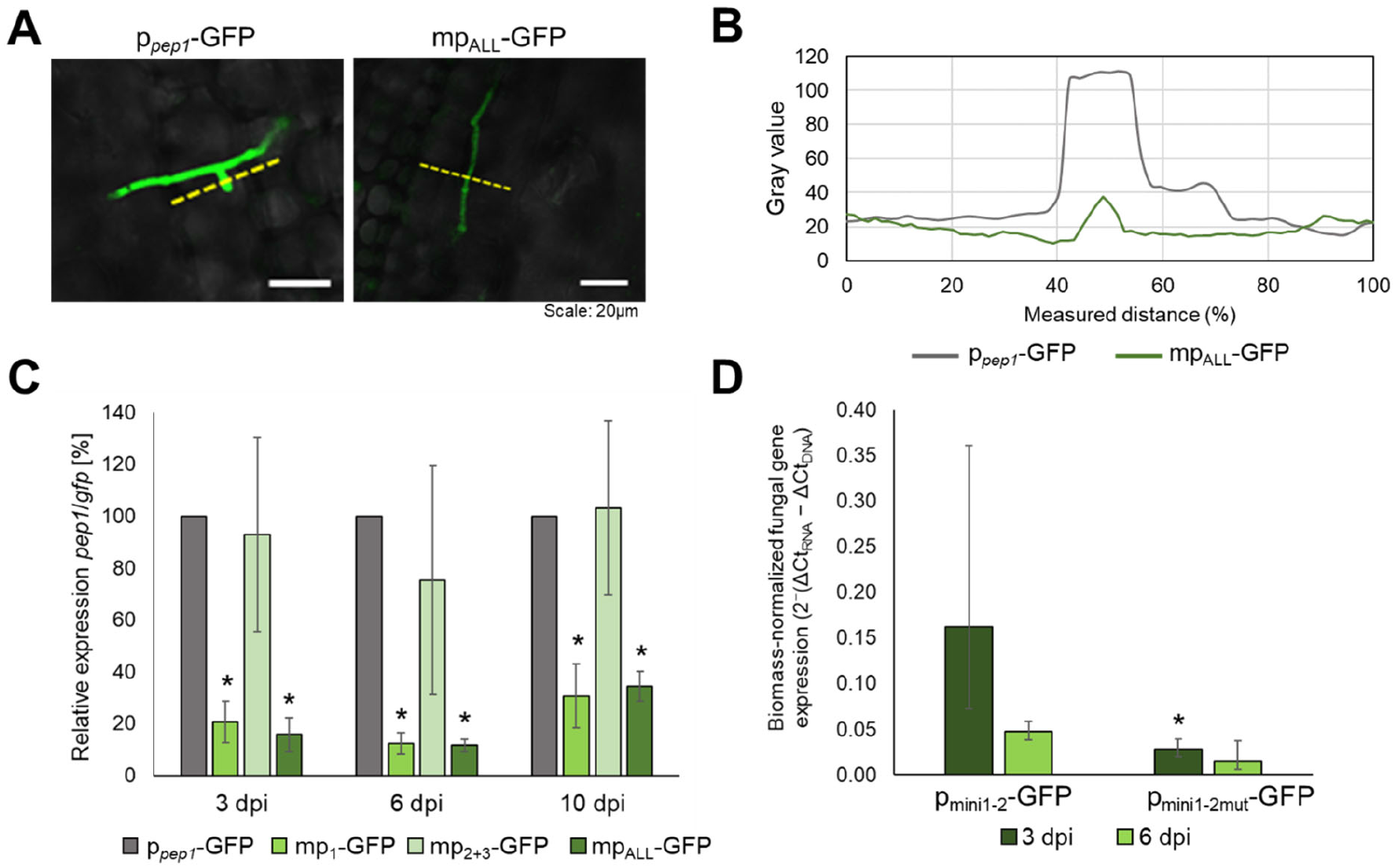
Mutation of phase-specific motifs reduce downstream *gfp* expression. *gfp* was placed under the control of the *pep1* promoter and integrated into the *ip* locus of *U. maydis*. To this end, a 600–1000 bp region upstream of *pep1* was cloned to drive *gfp* expression (p_*pep1*_-GFP). Motifs corresponding to the different phases (Figure 1) were mutated individually (phase 1: mp_1_-GFP; phase 2+3: mp_2+3_-GFP) or collectively (mp_ALL_-GFP). **(A)** Confocal microscopy images of maize seedlings infected with p_*pep1*_-GFP or mp_ALL_-GFP strains at 2 dpi. The dashed yellow line indicates the region used for GFP intensity measurements. **(B)** GFP intensity profiles quantified along the dashed yellow line using ImageJ. **(C)** Relative *gfp* expression is significantly reduced at 3, 6, and 10 dpi upon mutation of the phase 1 motif or all motifs. *gfp* and *pep1* transcript levels were normalized to the housekeeping gene *ppi*. Subsequently, *gfp* expression from the *ip* locus was normalized to *pep1* expression from the native *pep1* locus. **(D)** Artificial promoter harboring phase 1 motif p_mini1-2_ or p_mini1-2mut_, respectively, separated by 70 bp spacer region, followed by a minimal promoter (Promega) and a 5’UTR drives *gfp* expression in *U. maydis ip* locus. *gfp* expression was measured via qPCR normalized to *ppi* of *U. maydis* and further normalized to the fungal biomass based on gDNA. Data represent the mean ± SEM. Statistical significance was determined using Student’s t-test (p < 0.05).

To determine whether the GTGGG motif is sufficient to drive stage-restricted expression, we generated synthetic minimal promoters containing five tandem GTGGG copies separated by spacer sequences (p_mini1-2_). These minimal promoters drove GFP expression specifically during early infection, whereas mutation of the motif (AATCA; p_mini2-1mut_) abolished promoter activity (**Figure 4C,D**). This was not observed at 6 dpi. Together, these results demonstrate that the GTGGG motif contributes directly to early-phase effector gene expression *in planta* and can function autonomously as a synthetic regulatory module.

## Discussion

Biotrophic infection requires not only an appropriate effector repertoire but also its pre*cis*e temporal coordination of gene expression. The infection cycle of *U. maydis* comprises distinct developmental transitions, of which each stage imposes different physiological and host-interaction demands, including immune suppression, metabolic manipulation, and tissue reprogramming. It is therefore plausible that the pathogen deploys its effectors in a temporally structured manner, which is also reflected in the previous finding that effector gene expression occurs in an organ- and cell-type-specific manner (Skibbe et al., 2010; Matei et al., 2018) In *U. maydis*, hierarchical transcription factor cascades orchestrating pathogenic development have been extensively characterized (Romeis et al., 2002; Heimel et al., 2010, Lanver et al., 2017, 2018; Djamei et al., 2023). However, the *cis*-regulatory logic that encodes phase-specific effector expression at the promoter level has remained largely unresolved.

Here, we identified infection-phase-associated promoter motifs, analyzed their positional distribution relative to transcription start sites, and provide functional evidence supporting the regulatory relevance of at least one element. Whether the phase-enriched motifs identified in this study represent direct binding sites of known regulators or define additional regulatory layers remains to be determined. Stage-structured effector deployment has also been observed in obligate biotrophic rust fungi, where transcriptomic studies revealed temporally distinct waves of effector expression associated with penetration and haustorial development (Cantu et al., 2011; Dobon et al., 2016). Reanalysis of available RNA-seq data (Lanver et al., 2018) revealed clearly defined phase-enriched effector clusters, consistent with previous descriptions of transcriptional waves during infection (Lanver et al., 2017, 2018; Djamei et al., 2023). Our identification of promoter motifs enriched within these phase-specific clusters suggests that temporal effector deployment is, at least in part, encoded at the *cis*-regulatory level. These findings support a model in which the effectome is modular not only in function but also in time. Comparative motif analysis across multiple biotrophic fungi revealed that the GTGGG motif is consistently enriched in smut effectome promoters but not in rusts or powdery mildews. This suggests that certain aspects of effector promoter architecture may be lineage-specific and therefore, regulatory innovations may contribute to clade-specific pathogenic strategies.

Certain motifs also displayed enrichment downstream of annotated TSS. This may reflect alternative transcription start sites, regulatory elements within 5′ untranslated regions, or limitations in current gene annotations. Further high-resolution TSS mapping would help clarify these observations. The 1kb promoter of *pep1* contains 6 instances of the GTGGG [P1-2], 2 of TTGNCG [P2-1] and 3 of TSTTTS [P3-3], which is in line with its expression from 12 hpi to 12 dpi biotrophic development (Lanver et al., 2018). Among the identified motifs, GTGGG [P1-2] showed the strongest correlation with Phase 1 expression. Promoter mutagenesis within the *pep1* promoter resulted in a significant reduction of *gfp* expression during early infection stages. Furthermore, synthetic minimal promoters containing tandem GTGGG elements were sufficient to drive early-stage expression, whereas mutated variants lost this activity. These results provide direct evidence that at least part of the early effector wave is encoded at the *cis*-regulatory level. Pep1 is a key virulence factor required for successful biotrophic colonization (Doehlemann et al., 2009), and its correct temporal expression is likely critical for host compatibility. Although the cognate transcription factor remains unclear, motif similarity analyses suggest potential recognition by C_2_H_2_-type regulators, consistent with the importance of zinc finger transcription factors during early pathogenic development (Flor-Parra et al., 2006; Heimel et al., 2010).

Early pathogenic development in *U. maydis* is initiated by activation of the *b* mating-type locus and formation of the bE/bW heterodimer (Gillissen et al., 1992; Kämper et al., 1995), which induces downstream regulators such as Rbf1 (Heimel et al., 2010). Additional transcription factors including Biz1, Hdp2, and Mzr1 contribute to execution of the early pathogenic program (Flor-Parra et al., 2006; Heimel et al., 2010; Zheng et al., 2008). Later stages of infection involve regulators such as Zfp1 and the WOPR transcription factor Ros1, which promotes sporogenesis and late effector expression while attenuating early programs (Tollot et al., 2016; Cheung et al., 2021). The temporal association of GTGGG activity with early infection stages suggests potential integration within this established regulatory cascade. Future identification of the responsible transcription factor will be essential to position this motif mechanistically within the hierarchy. Even though the identified motifs demonstrate clear phase specificity in p^*pep1*^-GFP, which was demonstrated for the P1-2 motif using an artificial promoter, the phase-specific expression of any random gene could not be directly correlated with the presence of the respective motif in its promoter. This suggests that, while these motifs are effective individually, an integrated understanding of the surrounding *cis*-elements is also crucial.

Several limitations should be acknowledged. First, the identified motifs are relatively short (6–7 bp), increasing the probability of random occurrence. However, their positional enrichment, correlation with phase-specific expression, and functional validation (for GTGGG) collectively argue against purely stochastic distribution. Second, functional validation was primarily performed using the *pep1* promoter, and broader generalization across additional effector genes remains to be tested. Third, the identity of the transcription factors binding these motifs remains unresolved.

Beyond mechanistic insight, the identification of phase-associated *cis*-regulatory elements provides practical tools for infection-stage-specific gene expression and provide invaluable blocks in the generation of synthetic promoters. Defined minimal promoters with predictable temporal activity enable controlled misexpression experiments, stage-specific reporter systems, and synthetic rewiring of fungal gene expression during host colonization. More broadly, our findings suggest that temporal coordination of effector deployment represents an underappreciated evolutionary and functional layer of pathogen adaptation. Dissecting these regulatory elements may prove to be as critical as elucidating effector function itself for a comprehensive understanding of biotrophic plant-microbe interactions.

## Material and methods

### Prediction of total effectomes

The genomes, including chromosome and protein fasta files as well as annotation (GFF3 or GTF) files of 9 smuts: *Ustilago maydis* (GCA_000328475.2), *U. bromivora* (GCA_900080155.1), *U. tritici* (GCA_022963115.1), *U. nuda* (GCA_022963125.1), *U. loliicola* (GCA_022963135.1), *U. hordei* (GCA_900519145.1), *Sporisorium reilianum* (GCA_000230245.1), *S. scitamineum* (GCA_900002365.1), and *Melanopsychium pennsylvanicum* (GCA_900208305.1); 5 rusts: *Puccinia sorghi* (GCA_001263375.1), *P. striiformis* f. sp. tritici (GCA_000223505.1), *P. triticina* (GCA_000151525.2), *P. graminis* (GCA_000149925.1) and *Melampsora larici-populina* (GCA_000204055.1); and 5 powdery-mildews: *Blumeria hordei* (GCA_900237765.1), *B. graminis* f. sp. tritici (GCA_000418435.1), *Erysiphe necator* (GCA_000798715.1), *E. neolycopersici* (GCA_003610855.1) and *Golovinomyces cichoracearum* (GCA_003611235.1) were used in this study. For the prediction of the effectome, the protein fasta files were firstly subjected to an analysis with Phobius for the filtering of proteins without transmembrane domains (Käll et al., 2004). Thereafter, SignalP 6.0 was used for the filtering of the proteins which contain a signaling peptide, using the slow mode for eukarya (Teufel et al., 2022). Those proteins at this point, were considered as total secretome. For the determination of the total effectome, the total secretome of each species was used in an analysis with EffectorP 3.0 (Sperschneider & Dodds, 2022).

### Determination of orthologous genes

For the determination of gene orthologs among the tested species, the Orthologous Matrix standalone toolkit (OMA 2.5.0) was used (Altenhoff et al., 2019; Train et al., 2017). For this, the protein fasta files from all abovementioned genomes were used. In addition, to have outgroups for fungi and smuts, the protein fasta files for *Saccharomyces cerevisiae* (GCA_000146045.2), named yeast, and *Testicularia cyperi* (GCA_003144125.1) were used, respectively. The following parameters were modified in the parameters.drw file: OutgroupSpecies:=[‘yeast’], UseOnlyOneSplicingVariant:=false.

### Phylogenetic tree construction

The estimated species tree generated by the OMA standalone toolkit was used for the generation of the phylogenetic tree in Figure 1B. The inferred species tree in newick format was introduced to iTOL for visualization using the unrooted mode.

### RNAseq data analysis

Paired-end RNAseq reads from Lanver et al. (2018) were quality trimmed using Trimmomatic, selecting for per-base quality score in the start and end of the reads ≥ 20. Subsequently, reads were mapped to the *U. maydis* genome (GCA_000328475.2) with HISAT2 in stranded mode (Kim et al., 2019). Thereafter, successfully mapped reads were sorted with SAMtools and counted with HTseq-count using intersection-nonempty mode for stranded reads (Anders et al., 2015; Danecek et al., 2021). Reads deriving from the samples of 0.5, 1 and 2 dpi were added together to generate the Phase 1 timepoint; reads from 4 and 6 dpi, gave the Phase 2 timepoint; and subsequently, reads from 8 and 12 dpi were merged to create the Phase 3 timepoint. The differential gene expression analysis was performed with edgeR (Robinson et al., 2009), using a quasi-likelihood negative binomial generalized log-linear model to fit the count data. Normalized counts were calculated as counts per million mapped reads. Genes with FDR < 0.05 were considered as significant differentially expressed genes. For heatmaps, log_10_ transformed normalized counts were used.

### Filtering of phase-specific effectors

For the determination of phase-specific effectors, the following criterium was used: If 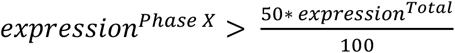, where *X* = 1, 2 or 3, is true, then the effector was considered as specific for Phase *X*. As expression, the normalized counts, in counts per million mapped reads, were used. The total expression (*expression*^*Total*^) was calculated as the sum of the normalized counts in all phases.

### Extraction of promoter sequences

The 500 or 1000 bp region upstream from the transcription start site was considered as promoter. For the extraction of promoter coordinates for the effector or background genes, the annotation files (GFF3 or GTF) along with the chromosome fasta files were used and the tool *flank* from bedtools was run with the following parameters -l 500 (or 1000 in case of 1000 bp promoter) -r 0 and -s, taking the gene orientation into consideration (Quinlan & Hall, 2010). Thereafter, the generated coordinates were used as input for the extraction of the promoter sequences in fasta format using bedtools *getfasta*.

### Motif enrichment and motif comparison

The 500 or 1000 bp promoter sequences were used for the motif enrichment analysis with the MEME Suite (Bailey et al., 2015). More specifically, the XSTREME tool was used with the following parameters: minimum motif width: 6, minimum motif width: 15, number of motif occurrences per sequence: any. For the motifs from the total effectome promoters, motifs with > 10% enriched sites and *P*-value < 0.05 were considered significant. For the significancy of phase-specific motifs, the thresholds for enriched sites > 55% and *P*-value < 0.05 were considered. Each significant motif was further manually curated to produce a consensus sequence. For the identification of phase-specifically enriched motifs, shuffled sequences were used as background, while for promoter motif enrichment of the total effectome, the secretome promoters of each tested species were selected as background. A further enrichment of each phase-specific motif versus the non-effectome background was estimated by counting the occurrences of the consensus motif in the effectome and non-effectome promoters of all tested species, followed by the calculation of the fold change enrichment.

The comparison between motifs was performed with STAMP using the Pearson correlation coefficient as comparison metric, the ungapped Smith-Waterman as alignment method, iterative refinement as the strategy for multiple alignment and tree construction with UPGMA (Mahony & Benos, 2007).Motif comparison to already existing databases was performed with the tool TomTom from the MEME Suite (Bailey et al., 2015). For this, the JASPAR CORE (2022) Fungi database was used. Comparisons with *P*-value < 0.05, and only the ones with good alignment of the core bases after manual observation, were considered good for the determination of the transcription factor groups which could be associated with the motif.

### Motif presence around the TSS

To count the presence of each phase-specific motif around the TSS of the *U. maydis* effectome and non-effectome, the HOMER Suite was used (Heinz et al., 2010). More specifically, the tool annotatePeaks.pl was deployed in *tss* mode, taking 1000 bp as a size around the TSS, to cover a 500 bp promoter region, and splitting the region into bins of 50 bp. The motif frequencies were then plotted for both effectome and non-effectome.

### Correlation between number of motifs and gene expression

In order to correlate the phase-specific gene expression potential of the *U. maydis* effectors with the occurrence of each putative phase-specific motif, at first working with one motif at a time, the effectors were split into groups based on the number of motifs present in their 500bp promoter, ranging between 0 and 6. Then, the average expression (normalized counts) for each phase, 1, 2 or 3, was calculated. Data were, afterwards, plotted generating a pattern of expression potential relative to the copies of the motif in the promoter. Significant differences were estimated by a Student’s t-test, with *p*-value < 0.05 cut-off. Promoter motif counts were correlated with gene expression levels using Spearman’s rank correlation coefficient. Scatter plots were visualized with slight horizontal jitter to reduce overplotting.

### Generation of mutants and subsequent infections

*Gfp* was cloned under the control of *pep1* promoter and integrated into the *ip* locus of *U. maydis*. Therefore, 1000 bp upstream of *pep1* were cloned to control *gfp* expression (pPEP1-GFP). The motifs of the different phases (Figure 1) were mutated separately (Phase 1: mp1-GFP, Phase 2:mp2-GFP) and all motifs were mutated (mpALL-GFP). Mutants harboring artificial promoters (miniP: derived from pGL4 (Promega, Madison, WI, USA) with Phase 1 motif and Phase 1 motif mutated driving *gfp* expression were cloned into the *ip* locus of *U. maydis*. Single integration of the constructs was verified by Southern blot and Sanger sequencing of the *ip* locus. 7-days old maize seedlings of the cultivar Golden Bantham were infected with pPEP1-GFP, mp1-GFP, mp2-GFP and mpALL-GFP as described previously (Redkar & Doehlemann, 2016).

### qRT-PCR and quantification of GFP expression

4-cm sections (1 cm below the infection site) of infected maize leaves were taken at 3 dpi, 6 dpi and 10 dpi. The frozen plant tissue was ground into a fine powder using liquid nitrogen followed by the extraction of total RNA using TRIzol (Thermo Fisher, Waltham, USA) according to the manufacturer’s protocol. Subsequently, a DNase I digest (Thermo Fisher) and cDNA synthesis (Thermo Fisher) was performed using the RevertAid H Minus First Strand cDNA Synthesis Kit, according to the manufacturer protocol. Relative expression of *gfp* was measured using the housekeeping gene *ppi* in a qRT-PCR. For quantification of *gfp* expression a normalization to the native *pep1* was performed. *gfp* expression in the *ip* locus was normalized to *pep1* expression in the native *pep1* locus, pPEP1-GFP was set to 100 and the ratio was calculated accordingly (Figure 5C). For the artificial promoters, *gfp* expression was measured relative to *ppi* and normalized on the fungal biomass using *ppi* of *U. maydis* and *gapdh* of maize. Due to the normalization method used, the standard deviation is displayed asymmetrically.

### Confocal microscopy and measurement of GFP intensity

To visualize GFP expression, the Leica TCS SP8 Confocal Laser Scanning Microscope (Leica, Bensheim, Germany) was employed. GFP was excited at 488 nm and detected within the 490-540 nm range. The microscopy images were processed using the Leica LAS X.Ink software. For quantification of GFP intensity, ImageJ was used.

## Acknowledgements

This project has received funding from the European Research Council (ERC) under the European Union’s Horizon 2020 research and innovation program (grant agreement no. 771035), as well as from the Cluster of Excellence on Plant Sciences (CEPLAS) funded under Germany’s Excellence Strategy—EXC 2048/1—project ID: 390686111. We thank Vanessa Volz and Niklas Gerling for generating *E. coli* constructs.

## Author Contribution Statement

Design of the project: GD, GS, JW; Bioinformatic analysis: GS: Experimental procedures: JW, KS, LH, NCS. Writing the manuscript: GS, JW with contributions from GD.

## Data availability

All source data are provided in this paper and its supplementary information. All microbial strains are available on request from the corresponding author.

## Supporting Information

**Figure S1:** Comparison of motifs enriched in the effectome promoters of biotrophic fungi.

**Figure S2:** Correlation between motif presence and gene expression potential.

**Table S1:** Details of the genomes of biotrophic fungi used in this study.

**Table S2:** Re-analyzed effector expression data from Lanver et al. (2018).

**Table S3:** Expression of *U. maydis* effectome.

**Table S4:** Estimation of the most probable transcription factor group associated with the phase-specific motifs.

**Table S5:** Confusion matrix.

**Table S6:** Sequence of Artificial promoters.

