## Supplemental Figures for "*Cis*-regulatory elements orchestrate phase-specific effector gene expression in *Ustilago maydis*"

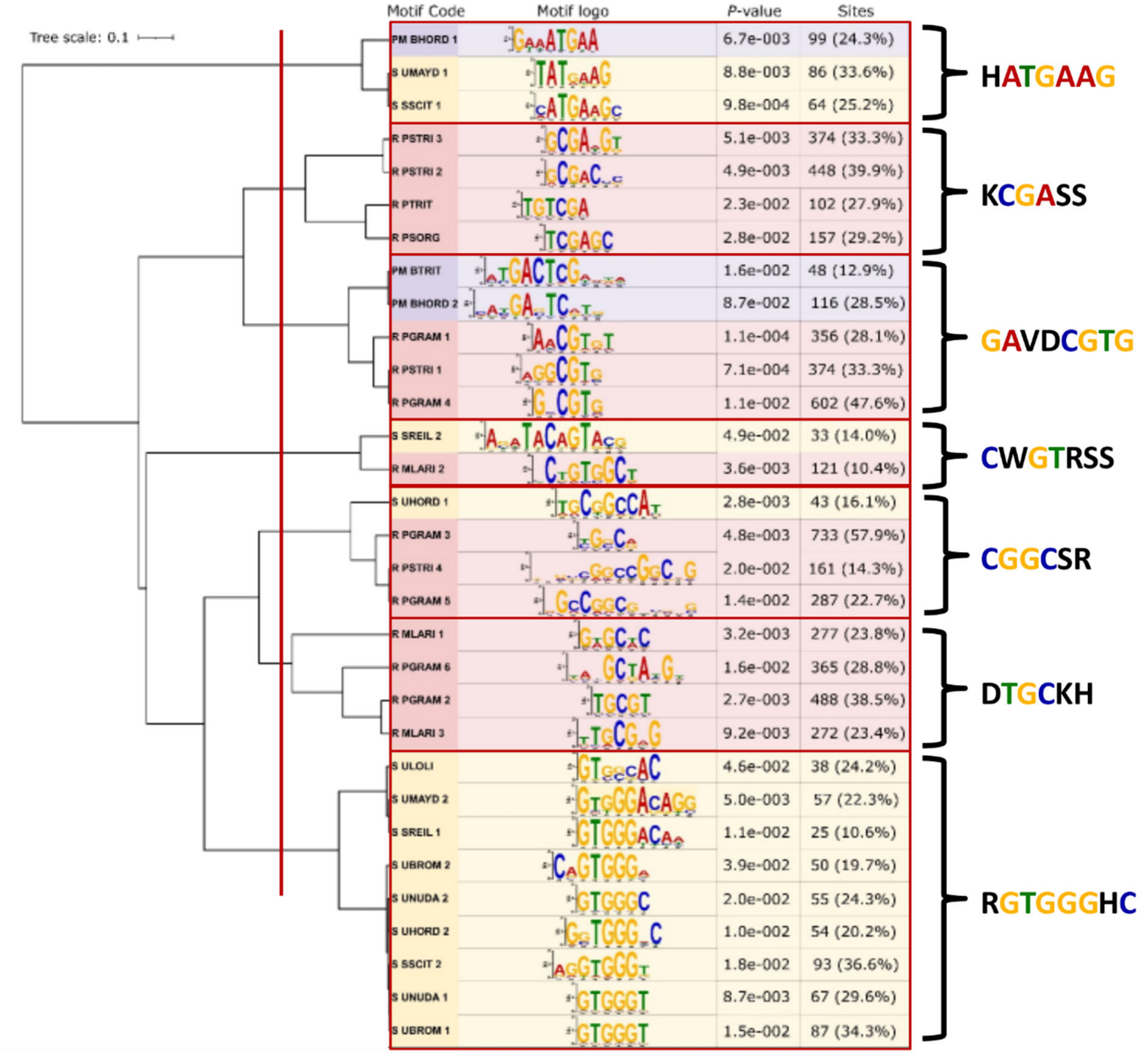


**Figure S1: Comparison of motifs enriched in the effectome promoters of biotrophic fungi.** The motifs were identified as enriched in the effectome promoters of the tested smuts, rusts and powdery-mildew species, with *p*-value < 0.05 and enrichment sites > 10%. Some species had more than one motif passing the set thresholds, which is reflected as a number at the motif code. Motifs were compared with STAMP and the UPGMA tree was generated with the same tool. The consensus core sequences of the groups generated after cutting the tree at the specified location (red line) were manually curated.

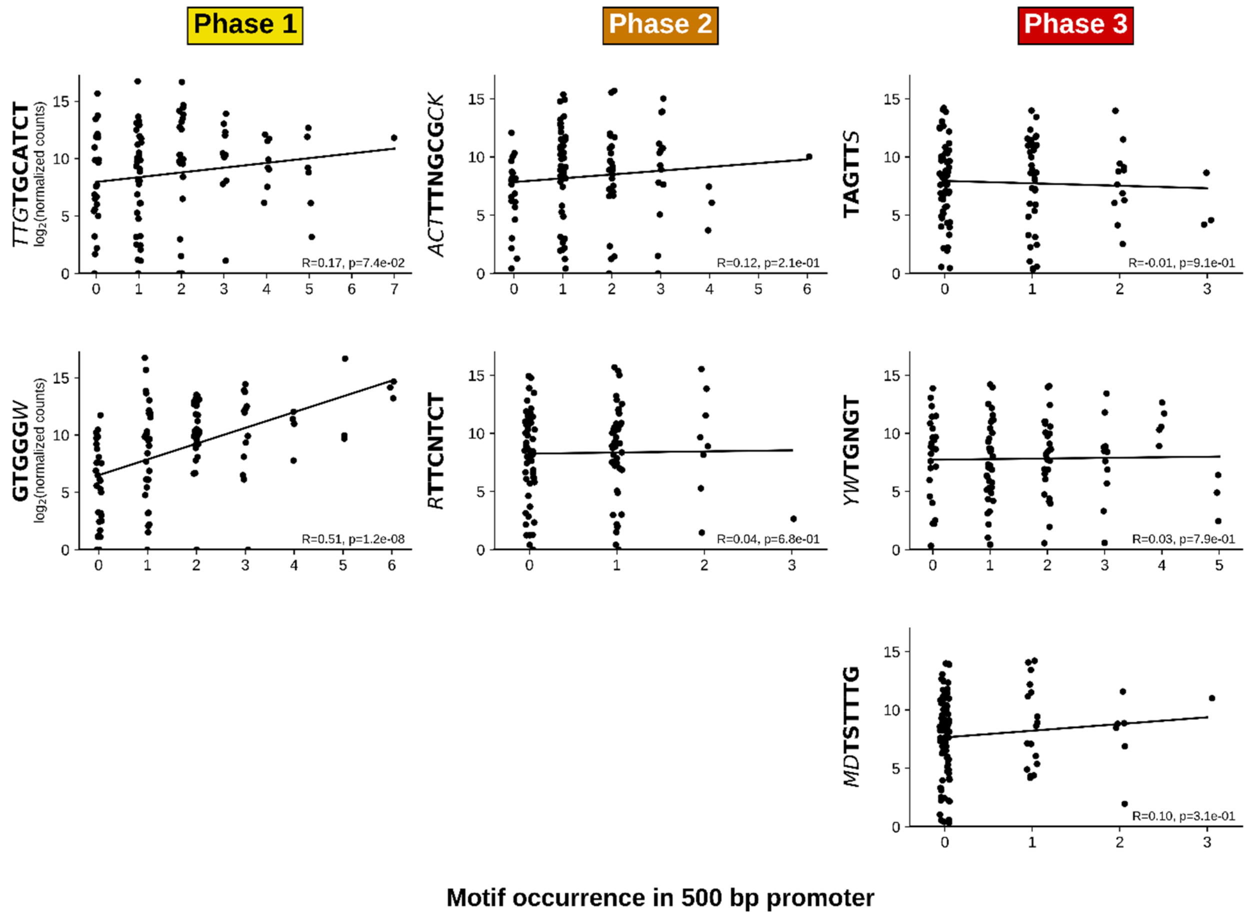


**Figure S2: Correlation between motif presence and gene expression potential.** Promoter motif counts were correlated with the maximum gene expression levels using Spearman's rank correlation coefficient. Gene expression values were log□-transformed (log□(x+1)). Scatter plots were visualized with slight horizontal jitter to reduce overplotting. Linear regression lines are shown for visualization purposes only. Motif cores were highlighted in bold within the full consensus sequence. The *R* statistic and *P*-value (p) of the correlation is depicted.
